# Impact of *Salmonella* genome rearrangement on gene expression

**DOI:** 10.1101/2022.05.04.490575

**Authors:** Emma V. Waters, Liam A. Tucker, Jana K. Ahmed, John Wain, Gemma C. Langridge

## Abstract

In addition to nucleotide variation, many bacteria also undergo changes at a much larger scale via rearrangement of their genome structure around long repeat sequences. These rearrangements result in genome fragments shifting position and/or orientation in the genome without necessarily affecting the underlying nucleotide sequence. To date, scalable techniques have not been applied to genome structure (GS) identification, so it remains unclear how extensive this variation is and the extent of its impact upon gene expression. However, the emergence of multiplexed, long-read sequencing overcomes the scale problem, as reads of several thousand bases are routinely produced that can span long repeat sequences to identify the flanking chromosomal DNA, allowing GS identification. Genome rearrangements were generated in *Salmonella enterica* serovar Typhi through long-term culture at ambient temperature. Colonies with rearrangements were identified via long-range PCR and subjected to long-read nanopore sequencing to confirm genome variation. Four rearrangements were investigated for differential gene expression using transcriptomics. All isolates with changes in genome arrangement relative to the parent strain were accompanied by changes in gene expression. Rearrangements with similar fragment movements demonstrated similar changes in gene expression. The most extreme rearrangement caused a large imbalance between the origin and terminus of replication and was associated with differential gene expression as a factor of distance moved towards or away from the origin of replication. Genome structure variation may provide a mechanism through which bacteria can quickly adapt to new environments and warrants routine assessment alongside traditional nucleotide level measures of variation.

## Introduction

Small nucleotide-level variations in bacterial genomes, such as single-nucleotide polymorphisms (SNPs), or small insertions and deletions (indels) can have huge effects, from altering antibiotic resistance to switching entire metabolic pathways on or off. Bacteria can also undergo changes at a much larger scale via chromosomal rearrangements, where large genome fragments shift position and orientation in the genome to ultimately produce different unique genome structures (GSs) without affecting the underlying nucleotide sequence. These large structural variations occur via homologous recombination around long repeat sequences, including transposases (Achaz et al. 2002), duplicated genes (Nakagawa et al. 2003), prophages (Brüssow et al. 2004; Fitzgerald et al. 2021), insertion sequence (IS) elements (Darling et al. 2008; Weigand et al. 2019, 2017; Lee et al. 2016) and ribosomal operons (Liu and Sanderson 1998; Page et al. 2020). Independent to the repeat sequence used as anchor points, large chromosomal rearrangements have been associated with speciation, diversification, outbreaks, immune evasion and host/environmental adaptation in bacteria (Hughes 2000; Fitzgerald et al. 2021; Brüssow et al. 2004). Such variation could offer several advantages for the survival of bacteria: it may rapidly provide varying phenotypes to enhance adaptability between different niches, it is reversible, and can alter expression patterns of many genes (Hughes 2000). Unlike other types of repeat sequences, ribosomal operons are present in all bacterial genomes and therefore genomic rearrangement is a mode of variation possible in all bacteria with two or more ribosomal operons (Page et al. 2020).

Short-read whole genome sequencing (SRS), alongside the ability to multiplex samples, has provided the necessary resolution and high-throughput required to regularly identify SNPs and other small nucleotide changes in bacterial species important in human health. However, whilst highly accurate, SRS reads are only hundreds of base pairs long and are therefore unable to resolve long repeat sequences to produce a complete assembly or detect genomic rearrangement. Historically, the detection of GS variation has been challenging and performed on an ad hoc basis with lower resolution methods such as long-range PCR or restriction enzyme digestion followed by pulsed-field gel electrophoresis (PFGE) (Liu and Sanderson 1996; Kothapalli et al. 2005; Matthews et al. 2011).

The emergence of long-read sequencing (LRS) technologies from Pacific Biosciences and Oxford Nanopore Technology (ONT) turns this situation around. LRS routinely produces reads of tens of thousands bases long, with potential to span across repeat sequences into the flanking DNA, producing complete assemblies that should ultimately allow the identification of GSs. The use of comparative genomic methods alongside visualisation programs has enabled multiple genomes to be aligned and compared which has helped highlight GS variation (Blom et al. 2016; Weigand et al. 2019; Fitzgerald et al. 2021; Darling et al. 2010) but investigating this variation using such methods is challenging to perform at high-throughput due to compute power requirements.

With more complete bacterial genomes being deposited into public databases, we previously demonstrated the ability to routinely identify GS variation from complete assemblies by developing a software tool called *socru* (Page et al. 2020). With *socru* we reported that many bacterial species important in human health display a wide range of GSs. The role GS variation plays in diseases may be underappreciated due to the lack of high-throughput methods required to routinely assess this variation.

Here we present the first use of LRS (via MinION, ONT) to confirm GSs originally identified by long-range PCR and show multiplexed LRS can be used to routinely monitor and determine GSs in a high-throughput manner. Our model system was *Salmonella enterica* serovar Typhi (*S*. Typhi), the causative agent of typhoid fever, a pathogen in which GS variation has been repeatedly observed (Liu and Sanderson 1996; Kothapalli et al. 2005; Liu and Sanderson 1998). *S*. Typhi appears particularly capable of producing different GSs (Liu and Sanderson 1998; Matthews et al. 2011); with more GSs found in *S*. Typhi than in all other *S. enterica* combined (Page et al. 2020). 45 GSs have been identified in *S*. Typhi via lab-based methods (Kothapalli et al. 2005; Matthews et al. 2011) and in 2019, we identified 17 GSs using *socru* from a total of 112 publicly available complete genomes (Page et al. 2020), 4 of which were novel. The ability to identify GSs in large numbers of bacterial genome sequences allows us to address the question of biological relevance of this, very common, form of bacterial variation.

Here we have used long-term *in vitro* culture of a laboratory strain to generate rearrangements, confirming these with long-range PCR and LRS. With these stable GS defined strains we investigated the impact of genome rearrangement on growth phenotype and gene expression.

## Results

### Laboratory-generated genome structure variation

After 4 months of long-term static culture at ambient temperature, different-sized individual colonies of the parent *S*. Typhi strain (WT) were observed, indicative of different growth phenotypes (Supplemental Fig. S1A). Both large and small colonies were picked at random for analysis.

### Genome structure by long-range PCR

To determine GSs of *S*. Typhi colonies via long-range PCR, 14 forward and reverse primers were designed (Supplemental Table S1) to bind to regions 100-900 bp downstream of the *rrs* gene and upstream of the *rrf* gene of each of the seven *rrn* operons, respectively. These primers were used to perform 91 individual long-range PCRs to test all possible combinations of neighbouring fragments. Primer combinations which amplified across an entire *rrn* operon produced a ∼6 kb band. The presence of seven different PCR products of correct sizes (∼6 kb) confirmed WT derivatives had seven genomic fragments and allowed their GSs to be determined (Fig. 1, Table 1 and Supplemental Fig. S2). WT itself was derived from Ty2 (see Methods) and confirmed to have the same GS 2.66 (17′35642′) (genome accession GCF_000007545.1.) (Deng et al. 2003).

**Figure 1.**
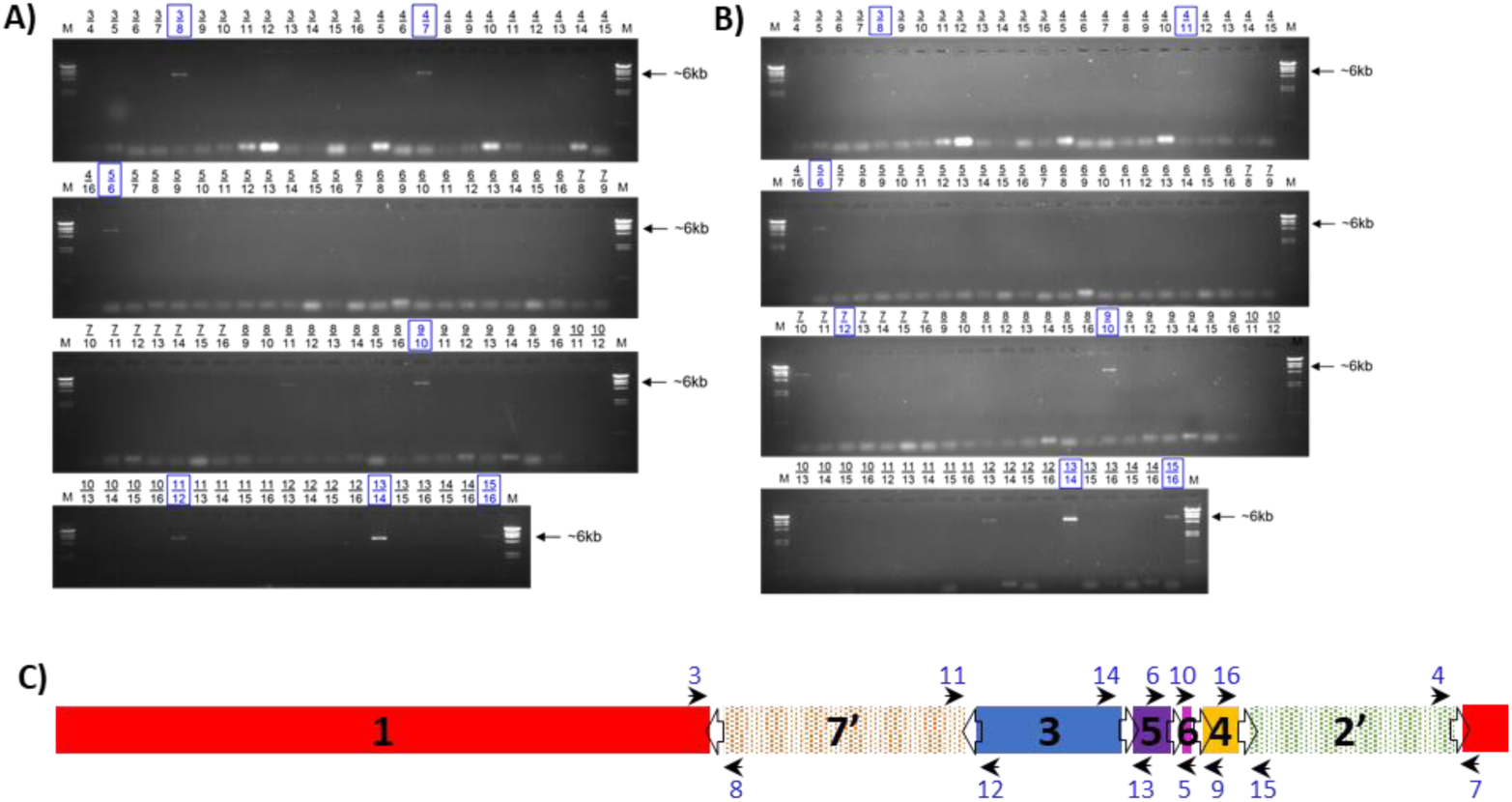
Long-range PCR for genome structure determination. Gel images of long-range PCR products of WT derivatives 7 (*A*) and T (*B*). Primer combinations are given above every well. Combinations indicated in blue boxes lead to the conclusion of the respective GS for that isolate. (*C*) Illustration of the primer binding sites within the *Salmonella* genome (Ty2 (WT) GS2.66, 17′35642′). Open arrows indicate the *rrn* operons and their orientation; black arrows indicate the direction and location of the primers numbered in blue; black numbers denote genome fragments.

**Table 1.**
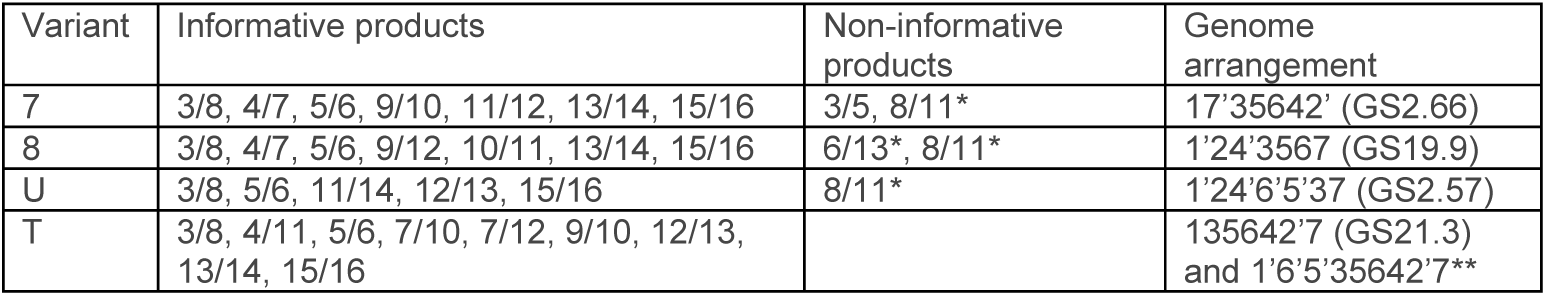
Arrangements determined by long-range PCR. Primer combinations resulting in PCR products that gave an informative 6 kb band were used to determine genome structures. Primer combination products were deemed non-informative either due to spurious bands of incorrect size or representing circularised fragments. *8/11 = circularised fragment 6, 6/13 = circularised fragment 5; **no GS assigned because the structure includes duplicated fragments.

Long-range PCR of variant 7 produced eight amplified PCR products of the correct size (Fig. 1A). The amplification of primer combination 8/11 represented the circularised fragment 6. The other seven bands indicated which fragments neighboured each other and demonstrated that isolate 7 maintained the parental GS described by 17′35642′ (GS 2.66, Fig. 2). Variant 8 produced nine amplified PCR products of 6 kb in length (Supplemental Fig. S2A) representing 1′24′3567 (GS19.9), where fragments 3, 5 and 6 had all undergone inversion in comparison to the parental GS (Fig. 2).

**Figure 2.**
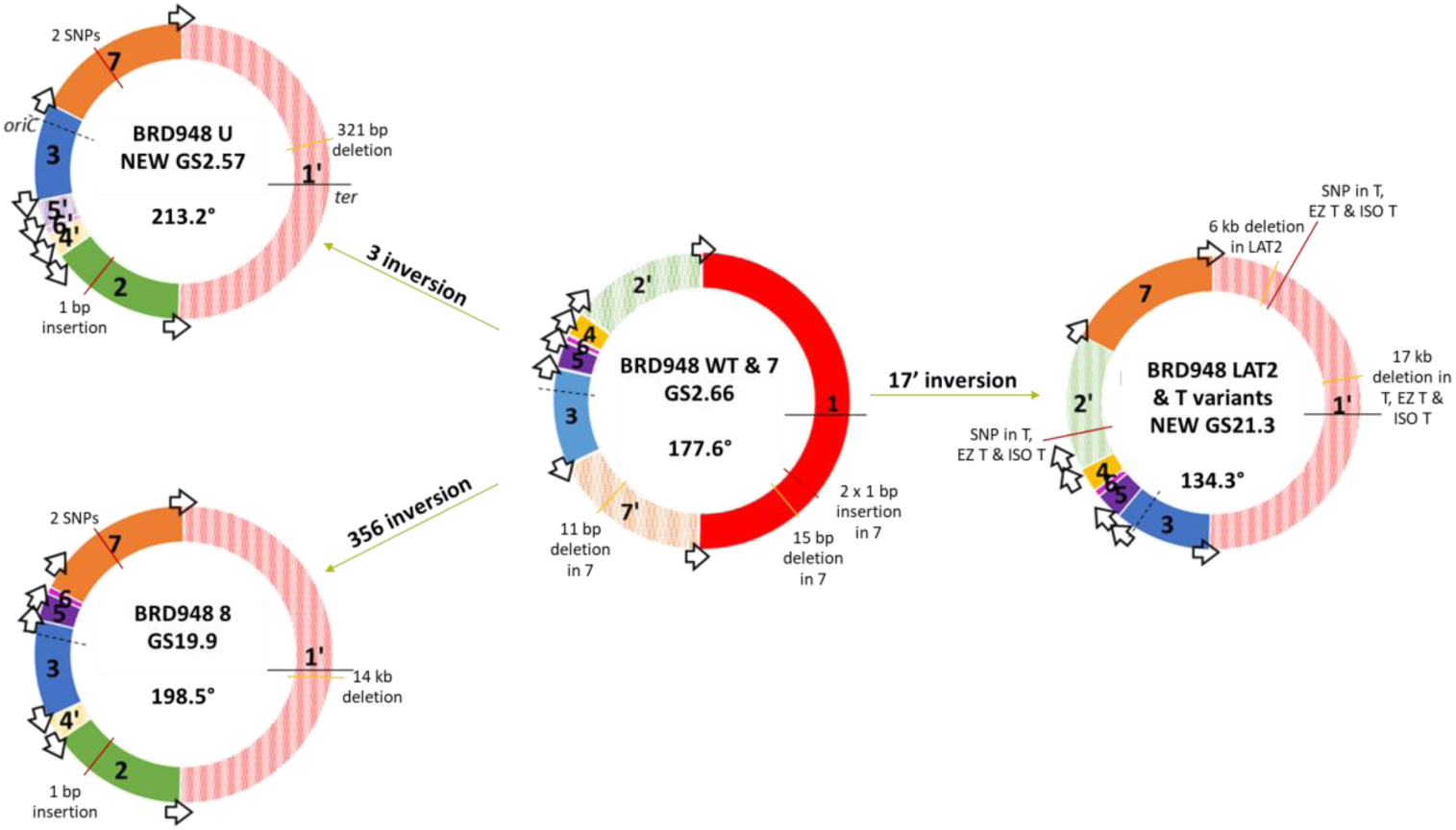
Genome rearrangements of variants relative to WT. Schematic showing variant genome structures (GSs) and the rearrangement of WT fragments required to achieve these. GS fragments are labelled in respect to the *Salmonella* enterica database reference LT2 (genome accession GCF_000006945.2) and drawn beginning with the largest fragment and working in a clockwise fashion around the chromosome. The fragment containing origin of replication (here fragment 3) has its orientation fixed to match the orientation of the database reference and therefore inversion of fragment 3 is depicted as the rest of the chromosome inverted. Inverted fragment orientations are denoted prime (‘) with striped colours. *Ori-ter* balance is given in degrees for each GS, going clockwise from *ter* to *oriC* as drawn. Arrows: ribosomal operons; *oriC* and dashed lines: origin of replication; and *ter* and black whole lines: terminus of replication. Data from genome sequencing used to identify insertions (red lines) and deletions (yellow lines) in each variant in comparison to WT; bp, base pairs.

Variant U produced five amplified PCR products of 6 kb in length (Supplemental Fig. S2B). These confirmed fragments 65′371′ [=1′735′6] and fragments 24′ were located together, respectively. Only one valid orientation existed for these two fragment blocks in relation to each other, as ribosomal operon direction must follow the direction of replication (Page et al. 2020). This gave the rearranged structure 1′24′6′5′37 (GS2.57), where fragment 3 has an inverted orientation in comparison to the parental GS (Fig. 2).

Long-range PCR of variant T produced nine amplified 6 kb PCR products (Fig. 1B). The informative bands indicated variant T displayed a potentially mixed GS population, with genome structures of both 135642′7 (GS21.3) and 1′6′5′35642′7 being present. These GSs both had fragments 1 and 7′ inverted relative to the parental GS (Fig. 2) and the latter had a duplication of fragments 5 and 6 within fragments 1 and 3.

These data confirmed the utility of long-range PCR in successfully identifying GS but also highlighted drawbacks for scalability e.g. requirement for 91 individual long-range PCRs (plus additional controls) to test all possible combinations of neighbouring fragments, and interpretation of the resulting gel to take into account informative versus spurious or misleading bands

### Genome structure by long-read sequencing

Following > 10 years of storage at -80 ^°^C, we re-cultured the parent strain and 4 variants, and performed LRS on these to determine if the GSs had remained stable and whether this method recapitulated the same GS as those identified by long-range PCR. For the parent strain and variants 7, 8 and U, DNA extraction was performed from over-night cultures. For variant T, overnight culture was repeatedly unsuccessful, and cells were instead harvested directly from an original glycerol stock prior to high molecular weight DNA extraction. We determined that 2.5×10^5^ cells/mL within this glycerol stock were still viable (Supplemental Fig. S3). Due to the limited amount of glycerol stock, trials at culturing this variant using alternative media (EZ-rich and iso-sensitest) were used to successfully revive T and make fresh glycerol stocks, named EZ T and ISO T respectively. For EZ T and ISO T, DNA extraction was performed accordingly from over-night cultures. All parent and variant DNA was sequenced on the MinION platform (ONT); long-read sequence data are presented in Table 2.

**Table 2.**
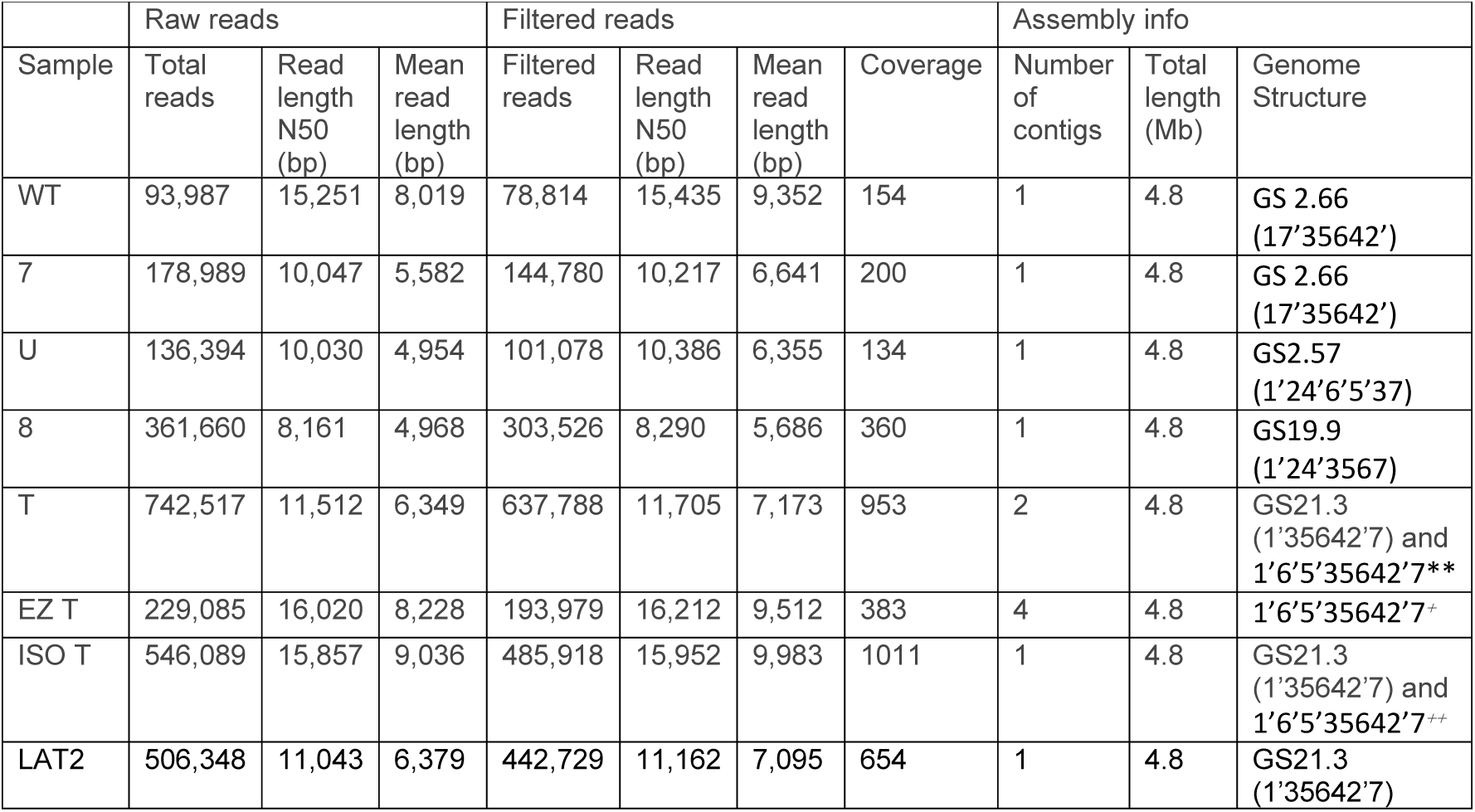
Long-read information from the WT parent strain and 7 derivatives. Filtered reads have length greater than 1 kb and min_mean_q of 50. Coverage based on length of Ty2 genome. **fragments 5 and 6 assembled on separate contig to rest of chromosome, mixed GS population; ^+^fragments 5 and 6 assembled on individual contigs, single GS population. ^++^all fragments assembled as single chromosomal contig, mixed GS population

Raw basecalled and demultiplexed fastq reads were filtered for high quality and for length greater than 1 kb. In our dataset, assemblies of the expected genome size were generated for all isolates. Genome structure assignments were determined from the assemblies using *socru* or prokka and Artemis Comparison Tool. Two isolates, U and T, have novel GSs not yet documented in the literature or public databases.

WT, 7, U and 8 each assembled into a single contig of ∼4.8 Mb which gave identical GSs to those determined by long-range PCR (Fig. 2, Table 2). In contrast, long-read assembly of T was in two contigs: 4.6 Mb (fragments 1′, 3, 4, 2′ and 7) and 0.2Mb (fragments 6 and 5) with the latter having twice the coverage of the former (Supplemental Fig. S4). A similar situation was seen with EZ T where fragments 5 and 6 were present on two individual contigs but still at twice the coverage of the main contig. To investigate the potential of a mixed GS population of 1′6′5′35642′7 and 1′35642′7 in these isolates, we searched the filtered reads for those which spanned fragments 3 and 5 and fragments 5′ and 3 (Supplemental Material, Supplemental Fig. S5, Supplemental Table S2). For T, the 3-5 bridge was present at approximately twice the presence of the 5′-3 bridge (208:111) indicating the two different GSs were present in roughly equal proportions and potentially explains why the assembly software struggled to either generate a complete assembly or assemble the dominant structure. For EZ T, the two bridges were present in approximately equal amounts (87:100), indicating the presence of 1′6′5′35642′7 only and the loss of GS21.3 from the population, in comparison to the original variant T. Assembly of ISO T gave a single contig of ∼4.8 Mb with a genome structure of 1′35642′7 (GS21.3). However, the two bridges were found in the filtered reads at a ratio of 2:1 (305:165), suggesting the presence of both GSs, as observed for T.

### Long-read sequencing as a method to monitor GS variation

Having confirmed LRS provided the same GS as long-range PCR, we used long-term culture in different media to generate genome rearrangements. Twelve large and small colonies were picked at random and processed for multiplexed LRS on a single MinION flowcell.

Sequencing was performed for up to 5 days to achieve the maximum amount of data for highest coverage, before data was demultiplexed and processed through our GS identification pipeline. Following LRS library preparation with the ONT rapid barcoding kit, assemblies of the expected genome size were generated for all tested colonies which had a mean read length of ∼10 kb and minimum ∼60x coverage. In one small, pin-prick colony (Supplemental Fig. S1B), LAT2, we observed genome rearrangement had occurred, producing a GS identical to isolate ISO T (1′35642′7, GS21.3) and was confirmed to contain only this GS via examination of filtered reads (Supplemental Table S2). The remaining colonies tested had not undergone rearrangement and had the parental GS.

### Nucleotide-level variation

Additional short-read whole genome sequencing was performed to generate hybrid assemblies for parent strain WT and variants 7, 8, U, T, EZ T, ISO T and LAT2. These gold-standard hybrid assemblies were evaluated with CheckM, which confirmed they were ≥ 99.66 % complete and contained <= 0.4 % contamination. The only exception to this was the completeness of T which was 93.07 %.

As expected, GS analysis of the hybrid assemblies gave identical results to those previously identified via long-read assemblies alone. Core genome SNP analysis of the variants confirmed that variants 7 and LAT2 were indistinguishable from the parent strain, WT. Isolates 8 and U were identical to each other but had 2 SNPs different to WT, at 4,629,839 bp (G→T) and 4,637,875 bp (C→A) in the Ty2 reference genome. T, EZ T and ISO T were identical to each other but harboured 2 different SNPs from WT: 677,285 bp (A→G) and 3,192,356 bp (C→T). All SNPs occurred in coding sequences, causing non-synonymous changes (Table 3). cgSNPs at 3,192,356 and 4,637,875 bp generated premature stop codons within the first and second domains of *tolC* and *treR* respectively. The SNP at 677,285 bp occurred in *rcsB* causing an amino acid located in the binding domain to change from a hydrophobic phenylalanine to a polar serine. The SNP at 4,629,839 bp occurs in t4482 (*licR*) and changes a negative charged aspartate to a large non-polar tyrosine.

**Table 3.**
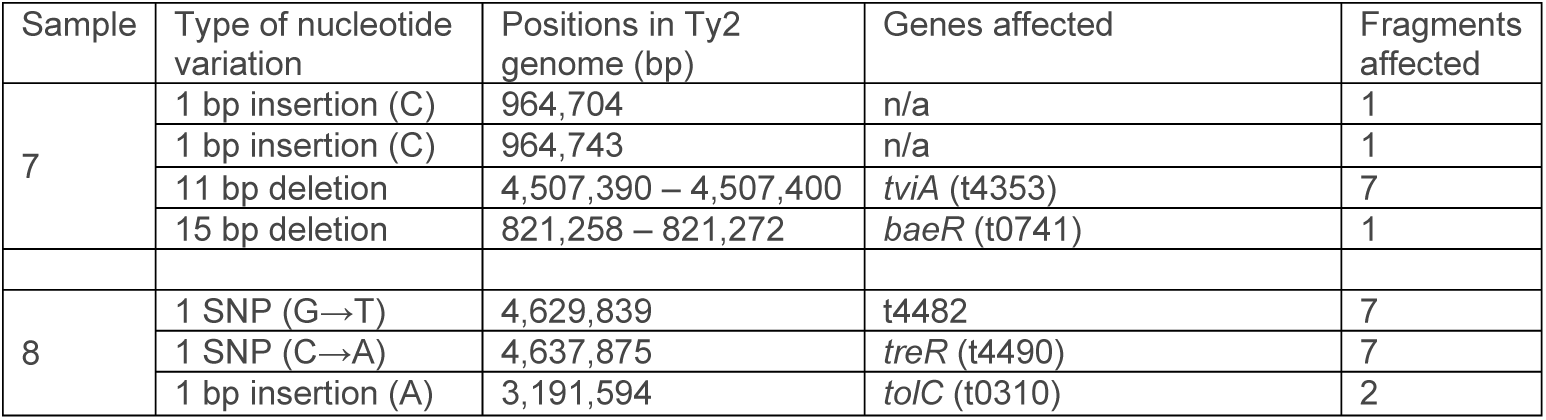
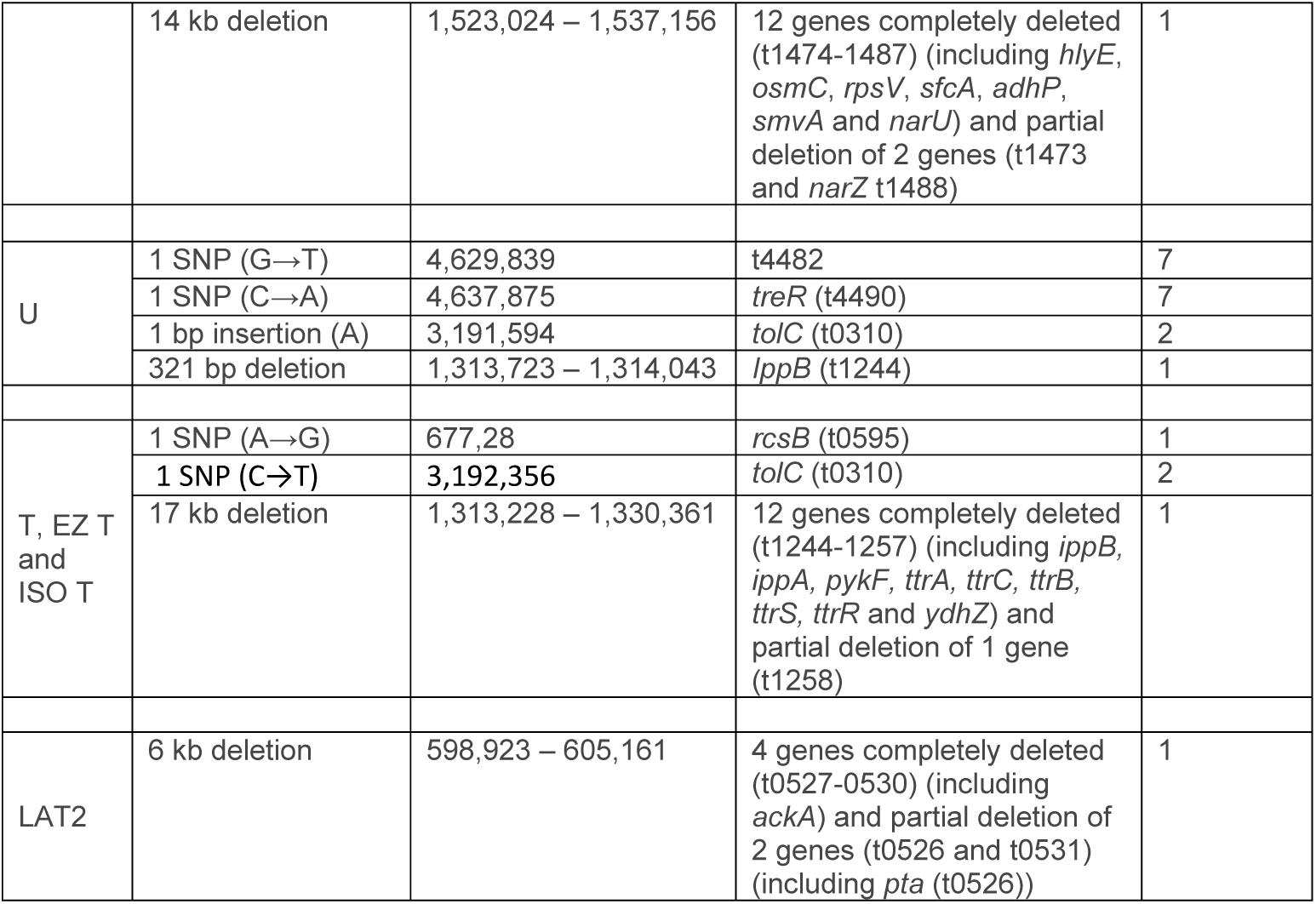
Nucleotide variation. SNPs, insertions and deletions identified in the 7 variants in comparison to the WT parent.

Further comparative genomics with Breseq revealed additional nucleotide variation, particularly associated with fragment 1 (Fig. 2, Table 3). Breseq was unable to detect the duplicated fragments 5 and 6 which are seen in T, EZ T and ISO T, as previously mentioned. Using the different levels of variation seen in the isolates generated in this work, we have generated the most parsimonious lineage (Supplemental Fig. S6).

### Impact of genome rearrangement on *ori-ter* balance

All rearranged isolates generated by long-term growth showed additional nucleotide level variation with all displaying indels and all but LAT2 having SNPs. In all cases, except isolate 7, the rearrangement caused the *ori-ter* balance to become more imbalanced. All the indels, except the smallest of 321 bp seen in U, occurred in the longer replichore which may represent some mechanism of compensation towards restoring *ori-ter* balance (Fig. 2). However, deletions ranged in size from 6-17 kb only resulted in shifting this balance by a maximum of 0.5°.

### Impact of genome rearrangement on growth rate

Variants 7, 8 and U showed similar growth phenotypes and colony sizes to the parent strain (Fig. 3, Supplemental Fig. S7). These growth phenotypes were consistent when repeated after 10 years in - 80 °C storage (Supplemental Fig. S7). From initial growth experiments, isolate T showed a clear reduction in colony size (Supplemental Fig. S1A) and growth rate compared to the parent.

**Figure 3.**
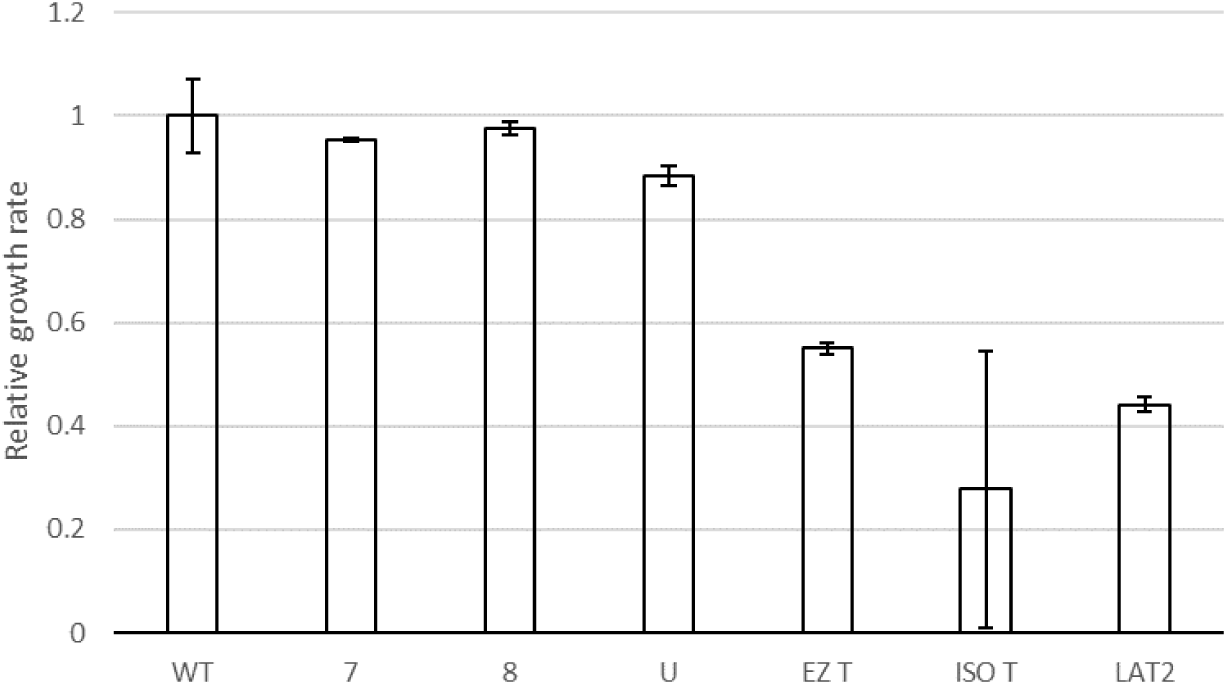
Growth rates of the 6 derivatives. Calculated for at least three independent biological replicates per isolate, relative to the WT parent strain. Error bars indicate standard deviation.

Subsequent growth experiments of revived T isolates and LAT2 showed similar reduction in colony size and growth rate (Fig. 3, Supplemental Fig. S1B).

### Impact of genome rearrangement on gene expression

The impact of rearrangement was explored using RNAseq to identify differentially expressed genes (DEGs). As T, EZ T, ISO T and LAT2 all had the same GS, with or without duplicated fragments, RNAseq was performed on WT parental strain and variants 7, 8, U and LAT2. Differential expression was determined for each variant in comparison to the parent strain, which harbours 4431 genes (Supplemental Table S3).

Isolate 7 (GS2.66) showed 63 significant DEGs (Supplemental Table S4). Only 13 genes were upregulated in isolate 7, which included the superoxide dismutase *sodA*, an indicator of oxidative stress also known to be positively regulated by BaeRS (Guerrero et al. 2013). This raises the possibility that the 15 bp in-frame lesion detected in *baeR*, whilst appearing within a response regulator receiver domain (Pfam: PF00072), may not have a functional impact on BaeR activity. The *cyo* genes encoding for the cytochrome *bo* (ubiquinol oxidase) terminal complex were also upregulated. Genes involved in Vi antigen (the capsular polysaccharide of *S*. Typhi which is a major virulence factor) and histidine biosynthesis were downregulated; the former may be partly due to the lesion detected within *tviA* which caused a frameshift mutation (Table 3).

For isolate 8 (GS19.9) and isolate U (GS2.57), 68 and 131 significant DEGs were identified respectively (Supplemental Table S4). Whilst representing different genome arrangements, they shared the same inversion of fragment 3, with the additional inversion of 5 and 6 in isolate 8 (Fig. 2). Not including the deletion of 14 genes in isolate 8, 83 % (45/54) of the significant DEGs in this isolate were also observed in isolate U. This included the upregulation of trehalose transport and utilisation (*treB, treC*) and of *ramA*, a transcriptional activator associated with multidrug resistance via AcrAB efflux (Nikaido et al. 2008), though no differential expression was observed for *acrAB* for either isolate. Tyrosine biosynthesis was downregulated in both (*tyrA*), as well as elements of glycolysis/gluconeogenesis (*pgk, eno*), with additional genes *pfkA, ppc* and *fba* downregulated in U.

By far the greatest impact upon expression was observed in LAT2, where 758 DEGs were identified (Supplemental Table S4). These were assessed in several ways: firstly, the genomic location of each significant DEG was plotted against the genome arrangement of both the parent (GS2.66) and LAT2 (GS21.3) (Fig. 4). This indicated that for LAT2, genes on fragment 1 between the terminus and fragment 3 appeared generally upregulated, coinciding with their shift of ∼ 800 kb towards the origin of replication. It also showed a general downregulation of genes on the other half of fragment 1 (between the terminus and fragment 7) in alignment with their shift of ∼ 800 kb away from the origin. Similarly, a general trend of downregulation was observed for fragment 7 genes, which had shifted ∼600 kb away from the origin.

**Figure 4.**
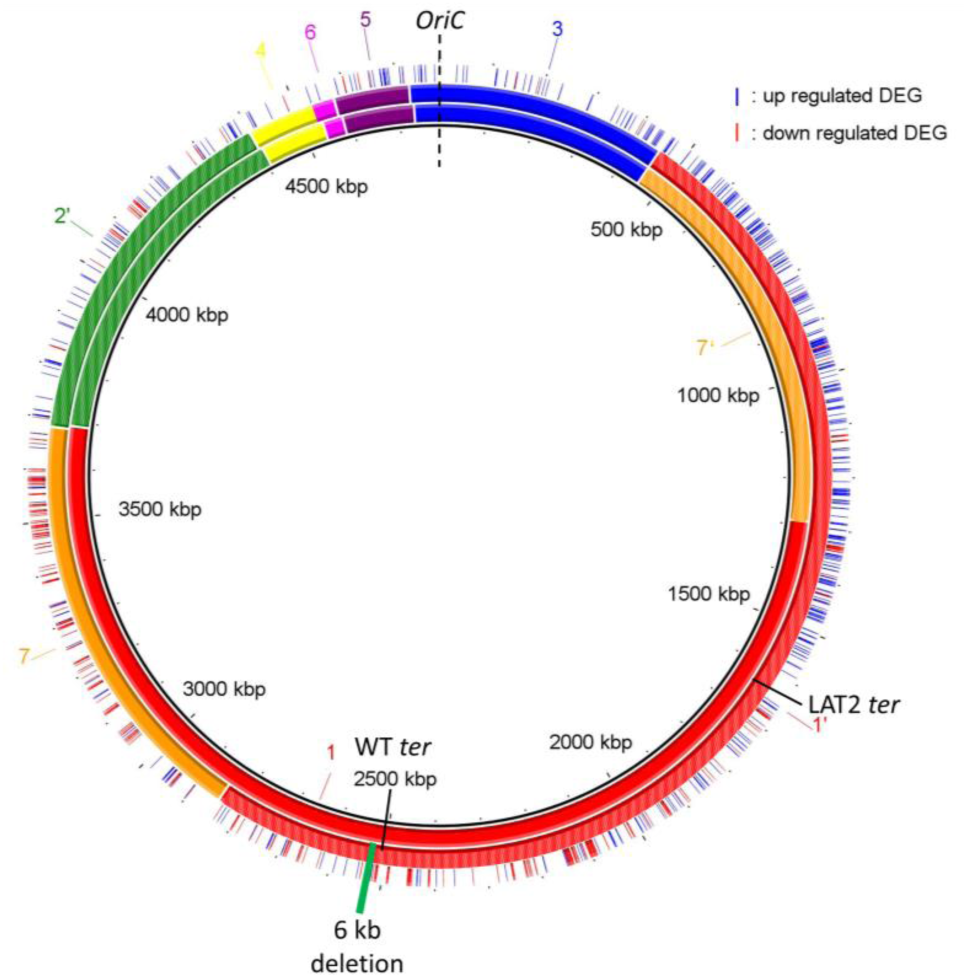
Gene expression in LAT2. BRIG representation of the WT genome (inner circle, GS 2.66 (17′35642′)) and the LAT2 genome (middle circle, GS21.3 (1′35642′7)). Genome fragments are numbered and shown as coloured blocks, inverted fragments are coloured with stripes (e.g. green fragment 2) as per (Page et al. 2020). Same origin (*oriC*, dashed black line) and different termini (*ter*, solid black lines) of replication are shown for each genome. Outer circle shows location of up (blue line) and down (red line) regulated differentially expressed genes (DEG). Deletion event denoted in LAT2 by solid green rectangle.

We therefore plotted genes per fragment by the distance they had shifted from the origin. This confirmed a large proportion of significant DEGs were found at the extreme ends of fragment 1 (Fig. 5A), though no strong correlation between direction of regulation and distance shifted to/from origin was observed across the fragment (R^2^ = 0.2836). For fragment 7, 81 % (99/122) of significant DEGs were downregulated (Fig. 5B).

**Figure 5.**
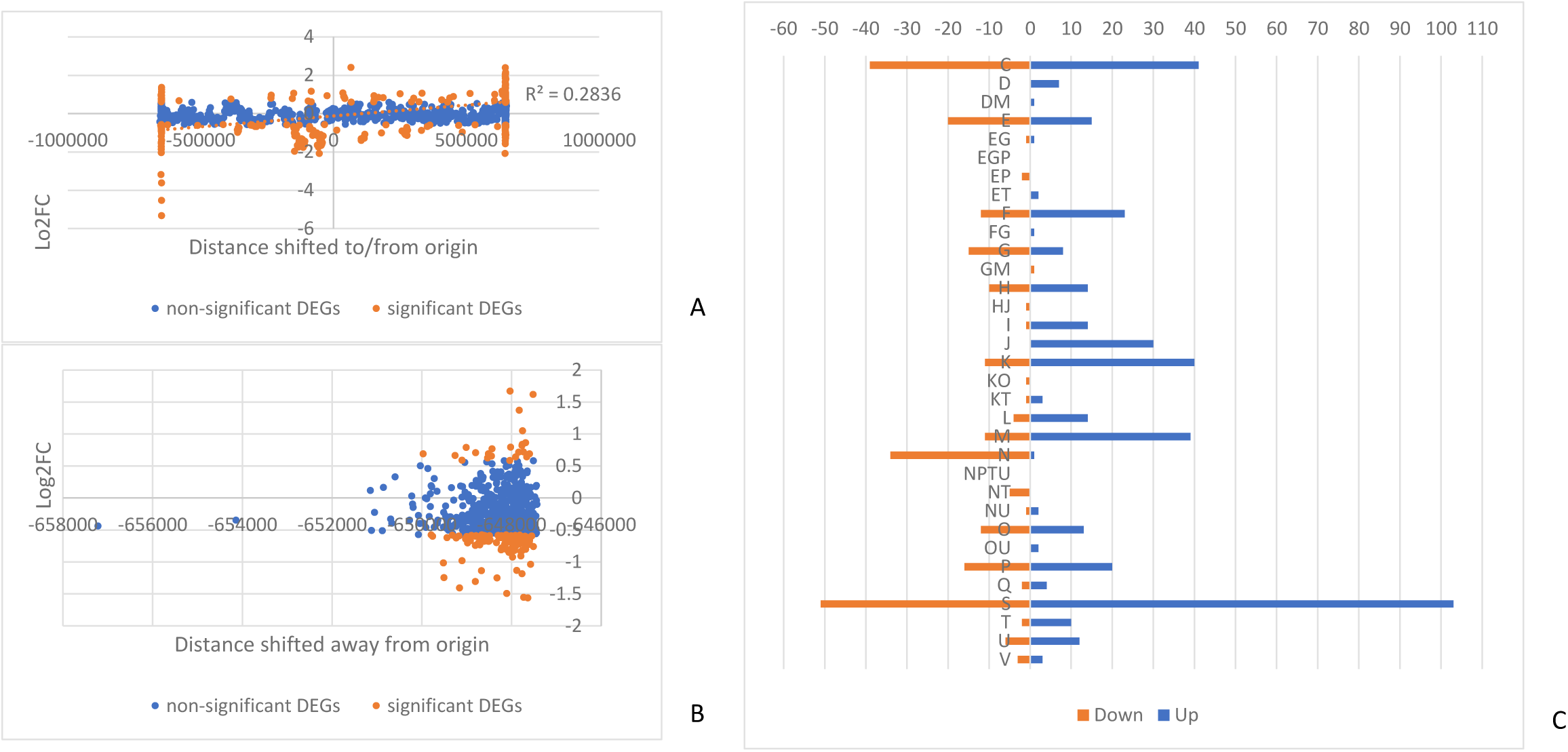
Impact of genome rearrangement on gene expression in LAT2. Graphical distribution of log2FC against distance a gene has moved towards or away from the origin of replication for LAT2 genes on (*A*) fragment 1 and (*B*) fragment 7. Genes coloured by non (blue) and -significance (orange). Linear correlation in (*A*) shown as orange dotted line. (*C*) Distribution of significant differentially expressed genes (DEGs) from LAT2 across COG categories. Down-regulated DEGs shown in orange, up-regulated in blue. COG categories: C Energy production and conversion; D Cell cycle control, cell division, chromosome partitioning; E Amino acid transport and metabolism; F Nucleotide transport and metabolism; G Carbohydrate transport and metabolism; H Coenzyme transport and metabolism; I Lipid transport and metabolism; J Translation, ribosomal structure and biogenesis; K Transcription; L Replication, recombination and repair; M Cell wall/membrane/envelope biogenesis; N Cell motility; O Posttranslational modification, protein turnover, chaperones; P Inorganic ion transport and metabolism; Q Secondary metabolites biosynthesis, transport and catabolism; S Function unknown; T Signal transduction mechanisms; U Intracellular trafficking, secretion, and vesicular transport; V Defense mechanisms.

We also investigated which clusters of orthologous gene (COG) functions were present in the significant DEGs (Fig. 5C). All but one of the genes (34/35) affecting cell motility (COG category N) were downregulated in LAT2. Conversely, genes in categories D – cell cycle control, I – lipid transport and metabolism and J – translation and ribosomal structure were almost all upregulated.

## Discussion

We have demonstrated that long-read sequencing can be used for GS identification, with the added benefit over long-range PCR of scalability alongside all the genetic information that comes from whole genome sequencing. We have shown that genome rearrangement has an impact on gene expression and growth rate with the greatest impact being when the *ori-ter* balance is most disturbed.

Considering that there are 1440 possible *S*. Typhi GS structures, it is of interest that the GS21.3 arrangement of T was recapitulated in an independent long-term growth experiment in isolate LAT2. This arrangement appears disadvantageous to bacterial growth due to the *ori-ter* balance being offset by ∼45° (Fig. 2), which was borne out in the growth rate analysis (Fig. 3). Theoretically, having fragments 1 and 3 next to each other in this arrangement of 1′35642′7 is the second most extreme *oriter* position that could be formed (the most extreme being where fragment 3 is also inverted: 1′3′5642′7). Even though isolates T and LAT2 were generated in different growth media, there are other conditions in common, including limited nutrients, growth waste products and anaerobic conditions. As such, we speculate that reduced growth rates seen in rearrangements such as GS21.3 may actually provide a selective advantage for survival in nutrient limited, or toxic, environments.

Given the growth effect of GS21.3, we investigated the impact that rearrangement had upon expression in all our GS arrangements. We observed that rearrangements with similar fragment movements demonstrated similar changes in gene expression. This was the case for isolates 8 and U which shared the inversion of fragment 3, and over 80 % of the DEGs in isolate 8 were also found in isolate U. In LAT2, GS21.3 caused a large imbalance between the origin and terminus of replication and was associated with differential gene expression as a factor of distance moved towards or away from the origin of replication, i.e. down regulation of most DEGs on fragment 7 and the greatest number of up/down regulated DEGs being found at the extremes of fragment 1. Specific COG function analysis highlighted that the metabolically costly production of flagella for cell motility was down regulated in LAT2 (> 30 genes all located on fragment 1), highlighting that a change in genome arrangement could be providing a mechanism of adaptation to poor nutrient levels.

In addition to GS changes, DNA sequencing also revealed SNP variation and larger deletions of hundreds to thousands of base pairs. The SNPs observed all caused non-synonymous changes and mostly occurred in outer membrane proteins. In isolate T (and derivatives EZ T and ISO T), one SNP results in a premature stop codon in the middle of TolC (Guan et al. 2015), a key outer membrane component of several multidrug efflux pumps. The second SNP was in *rcsB*, a transcriptional regulator that responds to cell envelope stress (Wall et al. 2018) and positively regulates Vi antigen biosynthesis (Virlogeux et al. 1996), caused a non-synonymous change within the DNA binding domain (Casino et al. 2018). However, since neither of these SNPs were found in LAT2, their effect on expression in this unbalanced arrangement will be the subject of future investigation.

The two SNPs (t4482, a putative *licR*-type regulator and *treR*) and indel shared by isolates 8 and U, suggest a link between genomic and genetic events. The SNP in *treR* resulted in a premature stop codon between its two protein domains (Hars et al. 1998). TreR negatively regulates *treBC* - aligning with the de-repression of *treBC* in these variants which have been shown in *E. coli* to have a role in mitigating against low osmolarity, by increasing conversion of trehalose to glucose via trehalose-6-phosphate (Vanaporn and Titball 2020). The indel was earlier in the *tolC* sequence than the SNP in the T variants, sending the sequence out of frame after 10 amino acids, resulting in a premature stop codon after 43 aa. As all upstream sequence remained unchanged, this did not affect *tolC* expression. However, the loss of TolC function in three variants with GS changes, by independent lesions in at least two, suggests that the export capacities of its associated pumps can be deleterious under the low-nutrient conditions used here.

In all rearranged isolates (U, 8, T and LAT2), deletions relative to the parent strain were identified in fragment 1 (Fig. 2). Strikingly, the largest deletions (14 kb and 17 kb) were very close to the terminus of replication in 8 and T, respectively. In *Salmonella enterica*, Koskiniemi et al demonstrated that deletion rates are highest near the terminus of replication and may be a mechanism to increase fitness in the particular conditions under which deletion occurs (Koskiniemi et al. 2012). This raises the possibility that genome rearrangement is a mechanism to target deletions.

To support investigation of GSs, long-read technology is key, and it is continually evolving. At the beginning of our routine monitoring of GSs, only 12-plex kits for the MinION were available to perform this work in a higher throughput manner. In 2020, to coincide with rapid large scale Covid sequencing, ONT released a 96-plex ligation kit which was quickly taken on by the community to sequence 96 samples containing 1 kb amplicons at once (Tyson et al. 2020). This throughput can now be leveraged to sequence up to 96 bacterial genomes per flowcell (Arredondo-Alonso et al. 2021), making routine GS identification the most accessible it’s ever been.

## Conclusion

In this study, we have identified 2 novel GSs, with one (GS21.3) being observed on two independent occasions. Through genomic and transcriptomic analysis, we have shown that the impact of rearrangement affects gene expression in similar ways across similar structural changes whilst the genome remains relatively balanced between the origin and terminus of replication, with more dramatic expression changes occurring in an unbalanced arrangement, accompanied by reduced growth rate. We also note that rearrangement appears to occur in conjunction with additional nucleotide variation, especially affecting gene presence near the terminus of replication. Incorporating routine identification of GS via long read sequencing will increase our understanding of the frequency of this type of variation and provide a strong foundation to systematically assess the role of rearrangement in bacterial adaptation.

## Methods

### Bacterial isolates included in this study

The *S*. Typhi strain used in these studies is WT, a long-term culture derivative of WT26 pHCM1 (Langridge et al. 2009). WT26 pHCM1 was originally derived from the attenuated Ty2-derived strain CVD908-*htrA*, which has deletion mutations in *aroC, aroD*, and *htrA* (Tacket et al. 1997), and further included a point mutation in *gyrA* and the multiple antibiotic resistance plasmid, pHCM1 (Turner et al. 2006). Long-term culture of WT26 lead to the loss of pHCM1 plasmid and the renaming of this strain to WT. Long-term, *in vitro* growth of WT in low salt LB (1 % tryptone, 0.5 % yeast, 0.5 % NaCl) generated 4 isolates (7, 8, U and T). After 10 years storage, isolate T was unable to be revived from glycerol stocks in original growth media and could only be revived using alternative media (EZ-rich (Teknova) and isosensitest (Oxoid)) which were used to make fresh glycerol stocks, named EZ T and ISO T respectively. Further long-term, *in vitro* growth of WT generated an isolate (LAT2) in isosensitest broth with a growth phenotype that deviated from that of the parent strain.

### Growth conditions for generation of different genome structures with long-term, *in vitro* growth

Long-term cultures were used to induce *in vitro* genomic rearrangement in *S*. Typhi WT. Due to the nature of attenuation in this strain, WT requires media to be supplemented with aromatic amino acid mixture (aro-mix) of L-phenylalanine, L-tryptophan, and L-tyrosine at a final concentration of 40 μg/mL and 2, 3-dihydroxybenzoic acid and ρ-aminobenzoic acid at a final concentration of 10 μg/mL.

Generation of variants 7, 8, U and T was achieved by growing a 50 mL aro-mix supplemented low salt LB culture of WT overnight at 37 °C, 180 rpm before leaving to grow at room temperature. After 4 months, 50 µL was plated out on low salt LB agar (Supplemental Fig. S1A), supplemented with aro-mix, and incubated at 37 °C for 48 hrs; individual colonies were picked for long-range PCR.

Generation of variant LAT2 and the other colonies tested by MinION sequencing was carried out as above and also extended to include aro-mix supplemented iso-sensitest media. Aliquots were plated out at intervals between 1 and 11 months; LAT2 was identified after 8 months of growth in iso-sensitest (Supplemental Fig. S1B).

### DNA extraction for long-range PCR

DNA extraction of *WT* derivatives was carried out using the Wizard Genomic DNA Purification kit (Promega). In brief, 1 mL of overnight *S*. Typhi culture, was harvested. Cells were pre-lysed in 600 µL of Nuclei Lysis Solution and incubated at 80 °C for 10 min. 3 µL of RNase A was added to the lysed cells and incubated for a further 15 min at 37 °C. 220 µL Protein Precipitation Solution was added to the lysed cells before being incubated on ice for 15 min. The precipitated protein was separated from the nucleic acids by centrifuged at 13.2 rpm for 15 min. 650 µL of the supernatant was mixed with 650 µL isopropanol before being centrifuged at 13.2 rpm for 15 min. The supernatant was discarded and the pellet was washed with 1 mL of 70 % ethanol, before being centrifuged at 13.2 rpm for 15 min. The supernatant was discarded and the pellet was left to dry. The dried pellet was resuspended in 45 µL of DNA rehydration solution.

### Long-range PCR for identification of genome structures

The primer sequences and combinations for detecting specific *rrn* (Supplemental Table S1) were designed using the program Primer3 Input 0.4.0 (http://frodo.wi.mit.edu/) and were synthesised by Sigma-Aldrich. All primers were aligned to the whole genome sequence of CT18 (Parkhill et al. 2001)to ensure specificity and no other matches with more than 80 % similarity were found. To ensure consideration of all options, every possible primer combination was used in 91 separate PCR reactions. PCRs were performed on 1 µL of DNA with 2X Fideli Taq PCR Master Mix (USB), 0.7 µM forward primer and 0.7 µM reserve primer in a total volume of 12.5 µL. The PCR conditions were: pre-incubation at 95 °C for 30 sec, amplification for 27 cycles at 95 °C for 25 sec, 59 °C for 1 min and 68 °C for 7 min, with a final extension at 68 °C for 7 min. Resulting *rrn* PCR products were separated out on 1 % agarose gels, before being detected using ethidium bromide staining (3 mg/mL).

### DNA extraction for sequencing

DNA extraction of *S*. Typhi isolates was carried out using a modified protocol of the PuriSpin Fire Monkey kit (RevoluGen). In brief, 1 mL of overnight *S*. Typhi culture, was harvested. Cells were pre-lysed in 100 µL of 3 mg/mL lysozyme, 1.2 % Triton X-100, and incubated at 37 °C, 180 rpm for 10 min. 300 µL lysis solution (LSDNA, RevoluGen) and 20 µL of 20 mg/mL Proteinase K (Qiagen) was added to the partly-lysed cells and incubated at 56 °C for 20 min. 10 µL of 20 µg/µL RNase A (Sigma) was added to the lysed cells and incubated for a further 10 min at 37 °C. 350 µL binding solution (BS, RevoluGen) and 400 µL 75 % isopropanol was added to the lysed cells before they were transferred to the spin column. Bound DNA was washed as per manufacturer’s instructions before being eluted in 2×100 µL of elution buffer (EB, RevoluGen) that had been pre-warmed at 65 °C. DNA concentration was determined using the broad range dsDNA assay kit (Thermo Fisher) on a Qubit 3.0 Fluorometer (Thermo Fisher). The quality of high-molecular weight DNA were assessed using the TapeStation 2200 (Agilent Technologies) automated electrophoresis platform with Genomic ScreenTape (Agilent Technologies) and a DNA ladder (200 to >60,000 bp, Agilent Technologies).

### Long-read sequencing

MinION libraries, containing 6/12 DNA samples, were prepared using the Rapid Barcoding Kit (SQK-RBK004, ONT) as per the manufacturer’s protocol. A pre-concentration step of 0.6x AMPure XP beads (Beckman Coulter) was performed on DNA samples which did not meet the manufacturer’s DNA input recommendations (400 ng in 7.5 µL). The library was loaded onto the flow cell according to the manufacturer’s instructions. Sequencing was performed on the MinION platform using R9.4 flow cells (FLO-MIN106, ONT) with a run time of up to 120 hrs. ONT MinKNOW software v1.4 was used to collect raw sequencing data and ONT Guppy v2.3.7 was used for local base-calling of the raw data after sequencing runs were completed. Python qcat command was used to de-multiplex samples.

### Short-read sequencing

Genomic DNA was normalised to 0.5 ng/µL with EB (10 mM Tris-HCl). 0.9 µL of TD Tagment DNA Buffer (Illumina Catalogue No. 15027866) was mixed with 0.09 µL TDE1, Tagment DNA Enzyme (Illumina Catalogue No. 15027865) and 2.01 µL PCR grade water in a master mix and 3 µL added to a chilled 96 well plate. 2 µL of normalised DNA (1 ng total) was pipette mixed with the 3 µL of the tagmentation mix and heated to 55 °C for 10 min in a PCR block. A PCR master mix was made up using 4 µL kapa2G buffer, 0.4 µL dNTPs, 0.08 µL Polymerase and 6.52 µL PCR grade water, contained in the Kap2G Robust PCR kit (Sigma Catalogue No. KK5005) per sample and 11 µL added to each well need to be used in a 96-well plate. 2 µL of each P7 and P5 of Nextera XT Index Kit v2 index primers (Illumina Catalogue No. FC-131-2001 to 2004) were added to each well. Finally, the 5 µL of Tagmentation mix was added and mixed. The PCR was run with 72 °C for 3 min, 95 °C for 1 min, 14 cycles of 95 °C for 10 s, 55 °C for 20 s and 72 °C for 3 min. Following the PCR reaction the libraries were quantified using the Quant-iT dsDNA Assay Kit, high sensitivity kit (Catalogue No. 10164582) and run on a FLUOstar Optima plate reader. Libraries were pooled following quantification in equal quantities. The final pool was double-SPRI size selected between 0.5 and 0.7X bead volumes using KAPA Pure Beads (Roche Catalogue No. 07983298001). The final pool was quantified on a Qubit 3.0 instrument and run on a High Sensitivity D1000 ScreenTape (Agilent Catalogue No. 5067-5579) using the Agilent Taestation 4200 to calculate the final library pool molarity.

The pool was run at a final concentration of 1.8 pM on an Illumina Nextseq500 instrument using a Mid Output Flowcell (NSQ® 500 Mid Output KT v2(300 CYS) Illumina Catalogue FC-404-2003) following the Illumina recommended denaturation and loading recommendations which included a 1 % PhiX spike in (PhiX Control v3 Illumina Catalogue FC-110-3001). Data was uploaded to Basespace (www.basespace.illumina.com) where the raw data was converted to 2 FASTQ files for each sample.

### Long-read and hybrid assemblies bioinformatics workflow

Bioinformatic analysis was performed on the open platform Galaxy. Prior to assembly, two steps were included to trim nanopore data. Filtlong v0.2.0 (https://github.com/rrwick/Filtlong) was used to trim nanopore data and only keep reads over 1 kb with a minimum mean quality score of 50. Porechop v0.2.3 (https://github.com/rrwick/Porechop) was used to remove sequencing adapters in the middle or the ends of each read. The long-read sequence correction and assembly tool Flye v2.5 (Kolmogorov et al. 2019) was used to assemble reads into contigs using an estimated genome size of 5 Mb. This long-read assembly was then polished with two rounds of Racon v1.3.1.1 (Vaser et al. 2017) and one round of Medaka v0.11.5 (ONT) using trimmed long-read data and corresponding overlapped reads generated by Minimap2 v2.12 (Li 2018). Hybrid assemblies were then generated by further polishing the final long-read assembly with two rounds of Pilon v1.20.1 (Walker et al. 2014) using short-read data and corresponding overlapped reads generated by Minimap2 v2.12 (Li 2018). Assemblies were evaluated for completeness and contamination with CheckM v1.0.11 (Parks et al. 2015).

GSs of isolates were then identified using two methods. Automatic identification of genome structure was performed by *socru* v2.2.2 (Page et al. 2020). Manual determination of genome order and fragment orientation was performed using Artemis Comparison Tool v18.0.2 (Carver et al. 2008) after annotation of the *rrn* operons with Prokka v1.14.5 (Seemann 2014). Within both methods, assembled genomic reads were aligned to the reference genome of *S*. Typhimurium LT2 which acted as a baseline for genome order and fragment orientation.

### Nucleotide variation analysis

Short-read data for WT and variants were analysed using the program breseq v0.24.0+2 (Deatherage and Barrick 2014), which outputs a list of probable mutations of various types and the sequence evidence for them. All analysis were run in consensus mode against the Ty2 reference sequence (RefSeq assession number NC_004631.1). Nucleotide variations which were common to WT and all variants, including deletions associated with the attenuation of WT strain, were not included in any further analysis. SNPs were checked using Snippy and Snippy-core v4.4.3 (https://github.com/tseemann/snippy). Large deletions (greater than 10 bp) were checked in Artemis Comparison Tool v18.0.2 (Carver et al. 2008).

### Original growth curve analysis of WT derivatives

Growth curves were generated by growing strains in triplicate in isosensitest broth at 37 °C with agitation. Overnight cultures were used to inoculate 200 µL isosensitest broth to an OD_600_ of ∼0.1 before OD readings were taken every 10 min over 11 hrs with a Fluostar Optima Microplate Reader (BMG Labtech).

### Repeated growth curve analysis of WT derivatives

Growth curves were generated by growing strains in triplicate in no salt LB broth at 37 °C with agitation. Overnight cultures were then standardised to an OD_600_ of ∼0.6, before a further 100X dilution was made. OD readings were taken every 15 min for 100 µL prepared cultures over 11 hrs with a Bioscreen C plate reader (Growth Curves Ltd). The growth rate was graphically determined by fitting a straight line on the exponential phase of the growth curve and calculating its slope.

### RNA extraction

RNA extraction of *S*. Typhi isolates was carried out, in triplicate for each isolate, using the All Prep DNA/RNA Mini extraction kit (Qiagen) following manufactures protocol. In brief, 100 µL of overnight culture was used to inoculate 10 mL EZ-media before being incubated at 37 °C, 180 rpm until an OD of ∼0.35-0.40 was reached (∼4 hrs). Cells were harvested by centrifugation at 4,000 g for 10 min and then resuspended in 100 µL RNAlater RNA stabilization reagent (Thermo Fisher). 600 µL buffer RLT Plus was added to the cell suspension before being pipetted mixed and transferred to an AllPrep RNA spin column. One volume (700 µL) of 70% ethanol was added to the flow-through before being pipette mixed and transferred to an AllPrep RNeasy spin. Bound RNA was washed as per manufacturer’s instructions before being eluted in 2×30 µL of RNAse-free water. RNA concentration was determined using the high sensitivity RNA assay kit (Thermo Fisher) on a Qubit 3.0 Fluorometer (Thermo Fisher). The quality of RNA were assessed using the TapeStation 2200 (Agilent Technologies) automated electrophoresis platform with RNA ScreenTape (Agilent Technologies) and a DNA ladder (50 to >6,000 bp, Agilent Technologies).

### RNAseq library preparation

From total RNA, the ribosomal RNA was depleted with the RiboCop rRNA Depletion Kit for Bacteria (Lexogen) using the Gram-negative (G-) probe mix according to the manufacturer’s protocol. RNAseq library preparation was carried out using a modified protocol of the QIAseq Stranded mRNA Select kit (Qiagen), which in brief used a fifth of the RNA input and reagents. The quality of RNAseq library were assessed using the TapeStation 2200 (Agilent Technologies) automated electrophoresis platform with D5000 ScreenTape (Agilent Technologies) and a DNA ladder (100 to 5,000 bp, Agilent Technologies). RNAseq librabries were sequenced on the Nextseq500 (Illumina) using a Mid Output Flowcell with the aim of obtaining 10 million reads per replicate (∼X2000 gene coverage). Data was uploaded to Basespace (www.basespace.illumina.com) where the raw data was converted to 2 FASTQ files for each sample.

### Differentially expressed gene analysis

Bioinformatic analysis was performed on the open platform Galaxy v19.05. The quality of raw sequences was ascertained using FastQC v0.72 (https://github.com/s-andrews/FastQC) before being quality control trimmed using fastp v0.19.5 (Chen et al. 2018). HISAT2 v2.1.0 (Kim et al. 2015) was used to align reads to the Ty2 reference sequence (RefSeq assession number NC_004631.1). Assignment of aligned reads to the genes of Ty2 was measured using featureCounts v1.6.3 (Liao et al. 2014) before DESeq2 v2.11.40.4 (Love et al. 2014), which is designed for the use with biological replicates, was used to determine differentially expressed genes from the count tables. The corrected p-value (p-adj), which is adjusted for multiple testing and controls the false discovery rate, was used to screen the DEGs. p-adj ≤ 0.05 was set as the threshold to judge significance of differential gene expression. After identifying significant DEGs, these were further screened using the absolute log2 fold change which was set to lLog2FCl ≥ 0.58, which is equivalent to lFCl ≥ 1.5, to judge the magnitude of the expression change.

Brig v0.95 (Alikhan et al. 2011) was used as a way to visualise the significant DEGs on a global scale using the parent WT genome, cut at dnaA to allow dnaA to be the beginning of the genome, as the backbone reference genome. The fragments of the parent and variate GSs were plotted as the first and second rings respectively to indicate the fragments involved in the genome rearrangement.

### PMA_xx_ real-time PCR bacterial viability test

A PMA_xx_ Real-Time PCR Bacterial Viability Test (Biotium Inc.), designed for selective detection of viable *S. enterica* cells in the presence of dead bacteria, was used to determine if any of the cells within a glycerol stock of isolate T were viable even though it was no longer culturable. See supplementary material.

## Supporting information

Supplemental Table S4: Significant differentially expressed genes

Supplemental Table S3: RNAseq

## Data Access

The Illumina and nanopore genome sequence data, RNA-seq data and hybrid assemblies generated in this study are available in DDBJ/ENA/GenBank databases under the Project accession number PRJEB52538 and per sample as: ERS11885537 (WT), ERS11885538 (7), ERS11885539 (8), ERS11885540 (U), ERS11885541 (T), ERS11885542 (ISO T), ERS11885543 (EZ T) and ERS11885544 (LAT2).

## Acknowledgments

The authors would like to thank Dave Baker and the QIB sequencing facility for support in Illumina DNA and RNA sequencing, Gemma Kay for advice in MinION setup and Satheesh Nair and Keith Turner for useful discussions on long range PCR.

EVW, JW and GCL gratefully acknowledge the support of the Biotechnology and Biological Sciences Research Council (BBSRC); this research was funded by the BBSRC Institute Strategic Programme Microbes in the Food Chain BB/R012504/1 and its constituent project BBS/E/F/000PR10349.

## Author contributions

EVW – Methodology, validation, investigation, formal analysis, visualisation, writing original draft, review and editing. LAT – Validation, investigation, formal analysis. JKA – Methodology, validation, investigation. JW – Conceptualisation, writing - review and editing. GCL – Conceptualisation, data curation, visualisation, writing original draft, review and editing.

## Supplemental Material

Supplemental Methods, Supplemental Figures S1-S7 and Supplemental Tables S1-S2

Supplemental Table S3: RNAseq

Supplemental Table S4: Significant differentially expressed genes

