## Supplemental Methods, Supplemental Figures S1-S7 and Supplemental Tables S1-S2 for "Impact of *Salmonella* genome rearrangement on gene expression"

Supplemental material

**A)**
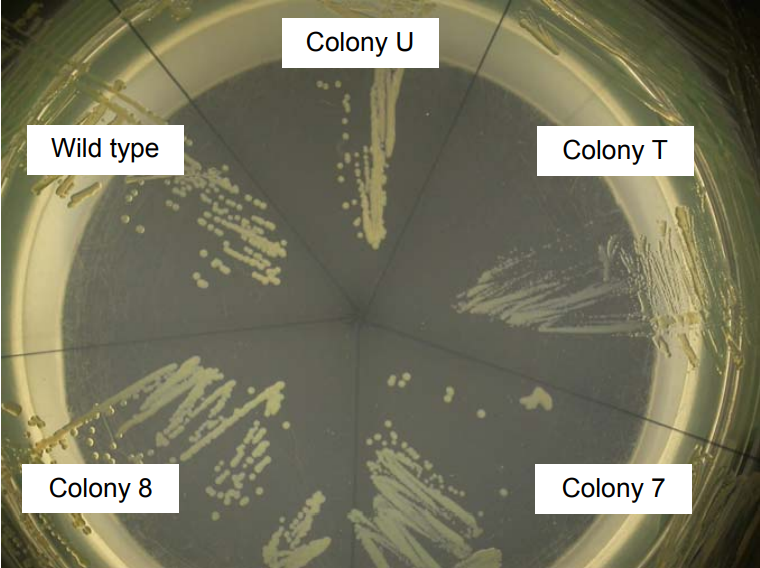
**B)**
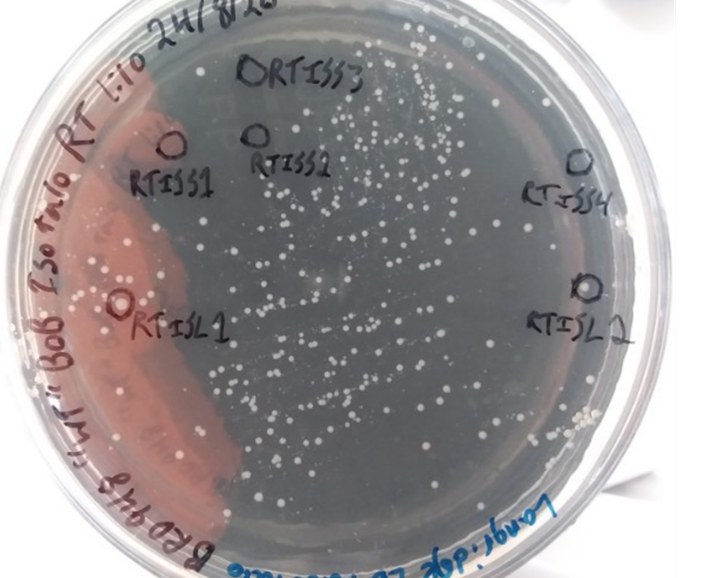

**Supplemental Figure S1.** Rearranged WT colonies. Plates of WT colonies showing different colony sizes after growth for 4 months in LB-NaCl broth (*A*) or 8 months in iso-sensitest broth (*B*). All cultures were supplemented with aro-mix and grown at room temperature. Pin-prick colony labelled RTISS4 contained isolate LAT2.

A)
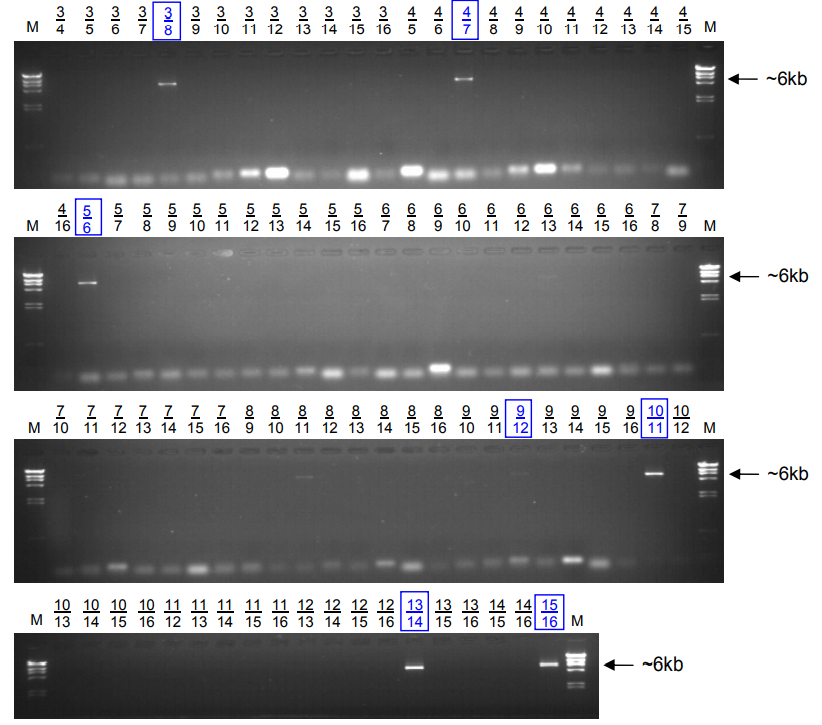
**B)**
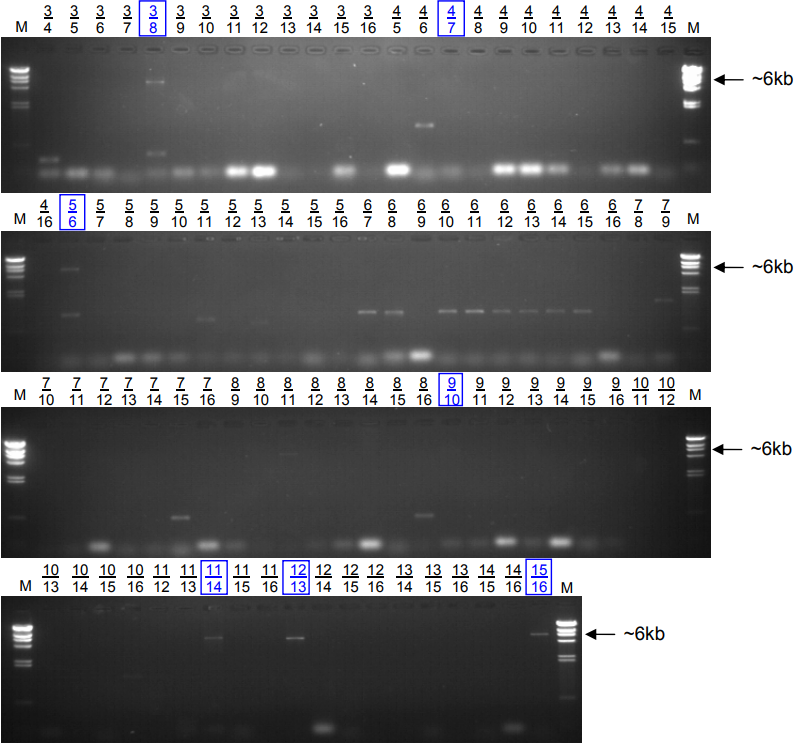

**Supplemental Figure S2.** Long-range PCR for genome structure determination. Gel images of long-range PCR products of WT derivatives 8 (*A*) and U (*B*). Primer combinations are given above every well. Combinations indicated in blue boxes lead to the conclusion of the respective GS for that isolate.

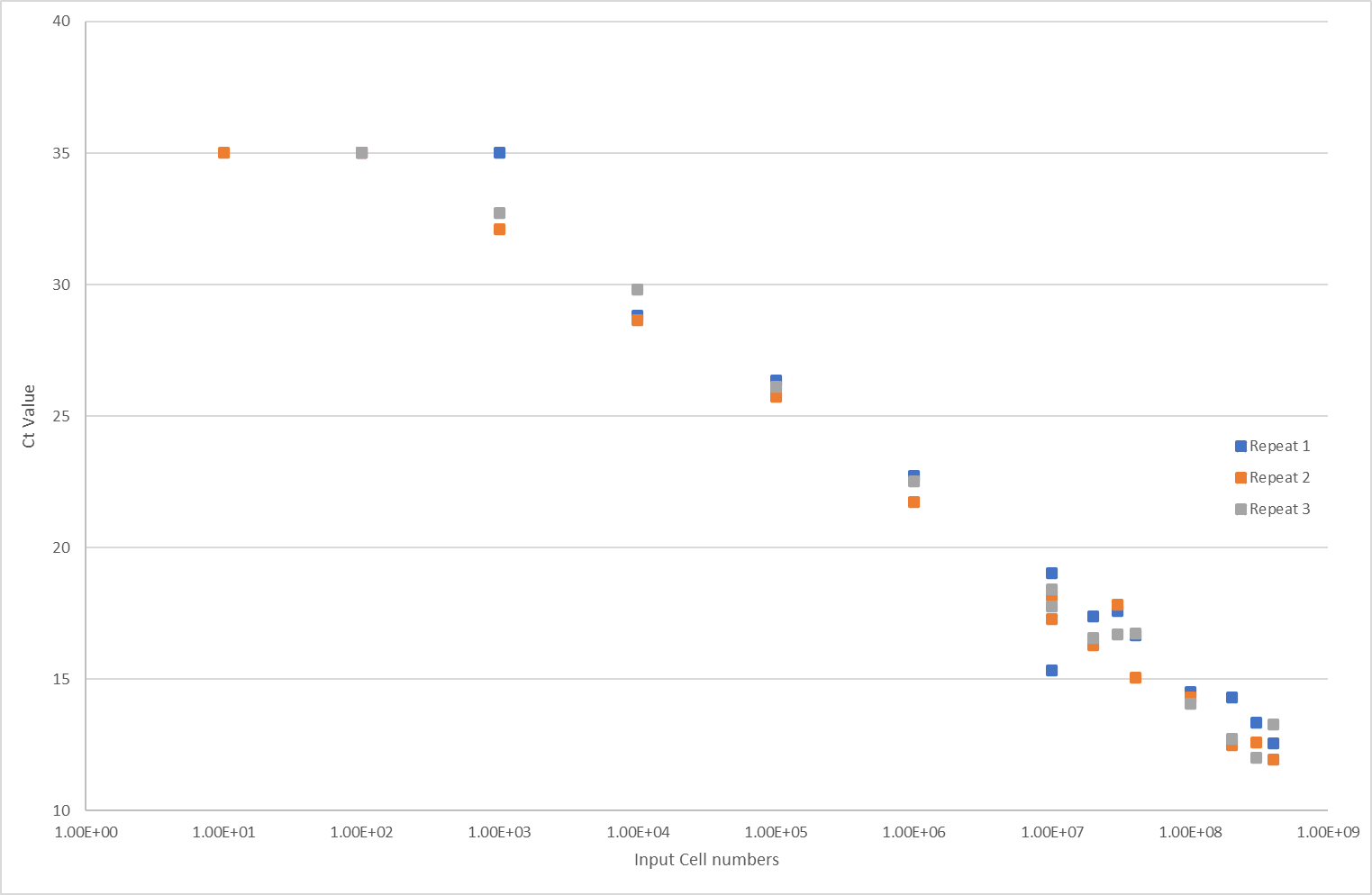

**Supplemental Figure S3.** PMAxx real-time PCR bacterial viability test. This is made up of three independent biological replicates at different cell inputs. With 400 μL T triplicate CT values of 26 were obtained which is equivalent to 100,000 alive cells, CFU of 2.5x10^5^/mL.

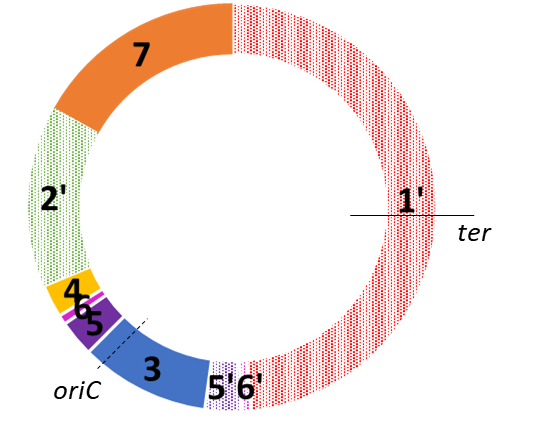

**Supplemental Figure S4.** Genome structure 1’6’5’35642’7. From isolate T which contain duplicated fragments 5 and 6.

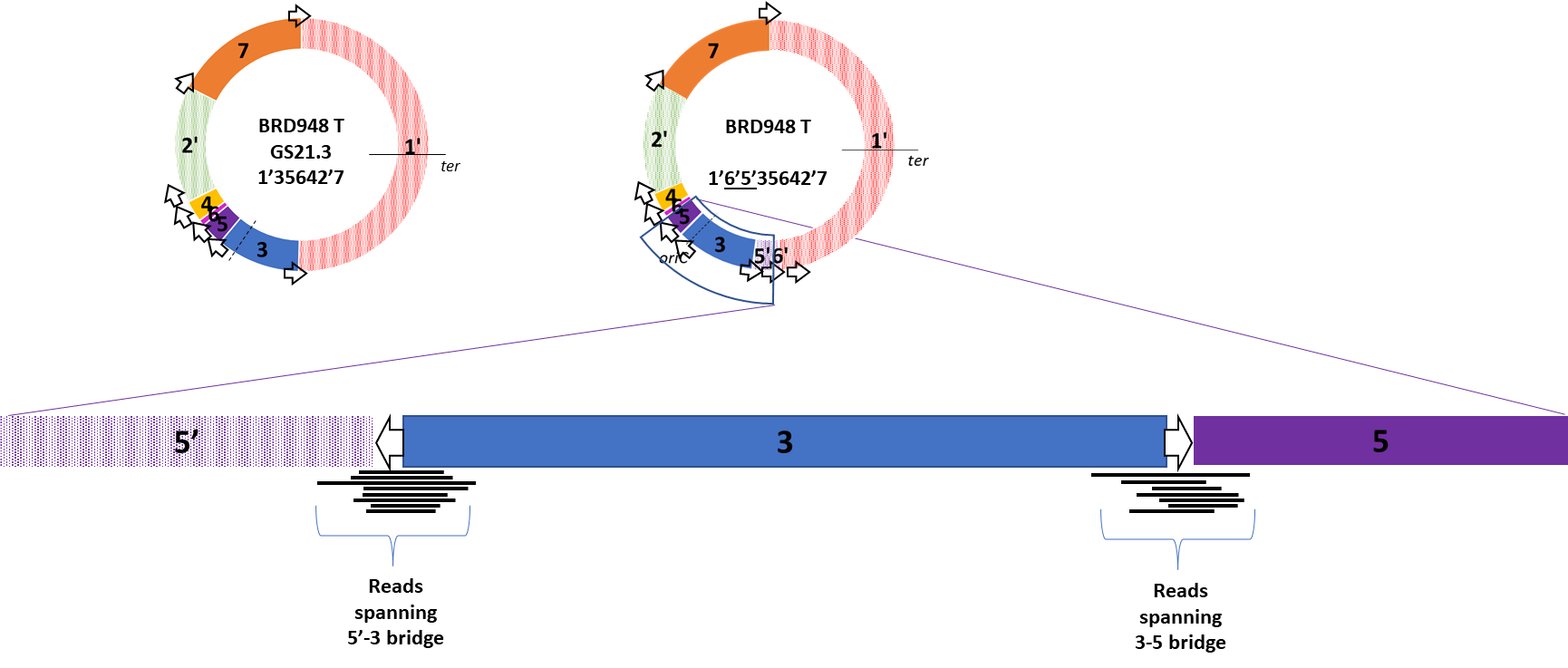
 **Supplemental Figure S5.** Schematic of the mixed GS population investigation. Above: T variant structures GS21.3 (1’35642’7) and 1’6’5’35642’7. Below: Expanded 5’->3-> 5 section demonstrates which reads were searched for, bridging fragments 3 and 5 (3-5) and fragments 5’ and 3 (5’-3).

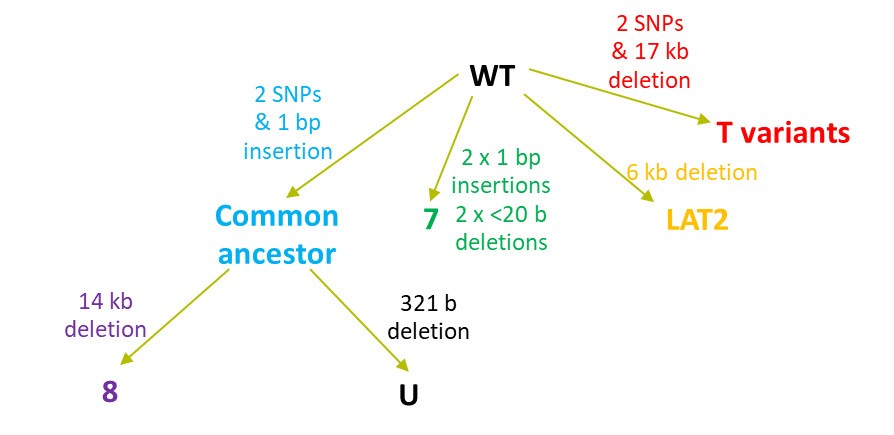

**Supplemental Figure S6.** Parsimonious lineage of isolates. Most parsimonious evolutionary trajectory leading from the parent strain, WT, to the variants described in the main text.

**A)**
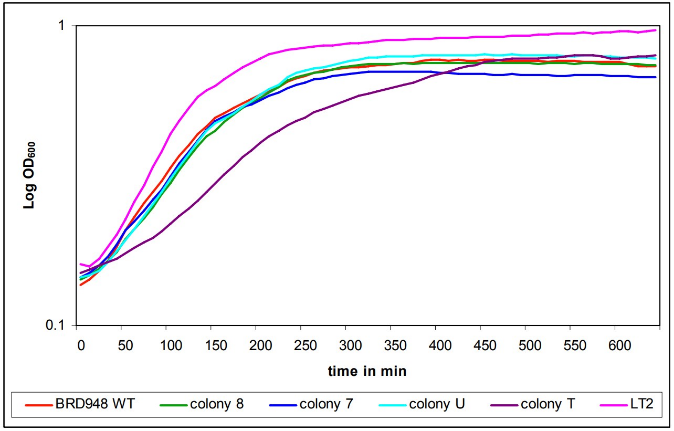
**B)**
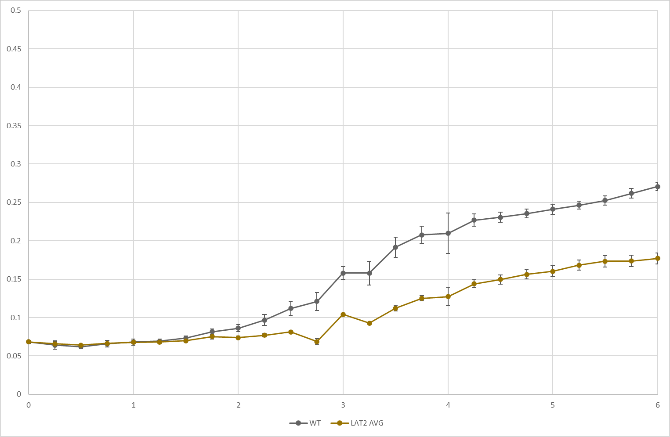

**C)**
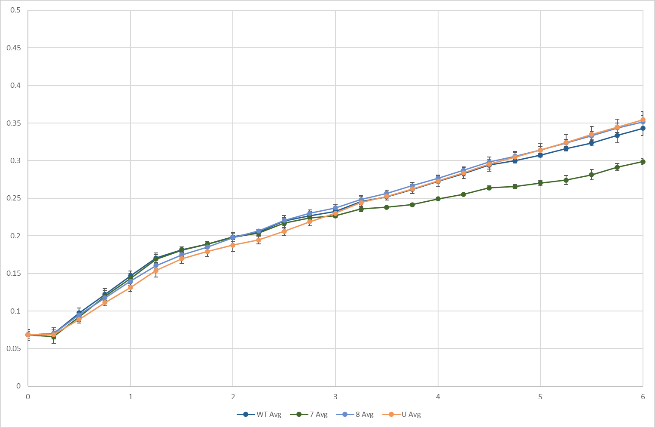
**D)**
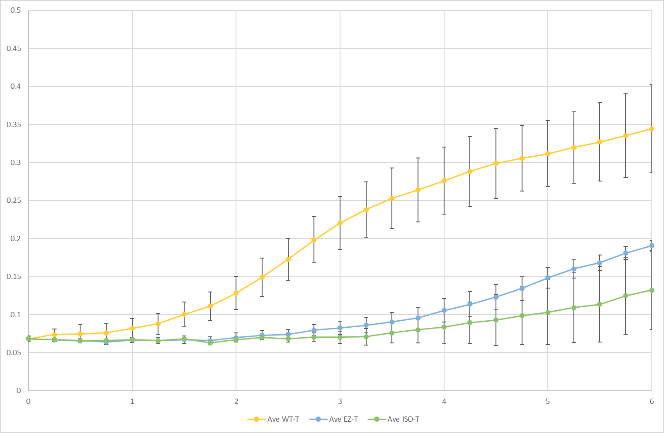

**Supplemental Figure S7.** Growth curves of BRD948 isolates. *A*) Growth curves performed over 10 years prior to repeated growth curves shown in *B*), *C*) and *D*). Dataset in A includes *Salmonella enterica* serovar Typhimurium LT2.

| Primer | Sequence (5’→3’) | Position in CT18 genome  (5’ end of primer) |
| --- | --- | --- |
| 3 | AAGCACGTGTAAAGGATAGTTCATC | 2711858 |
| 4 | AACCGTATTGATGAAGTTGTGGTAT | 2718308 |
| 5 | TTCATTTTTGAAGATACTGTCACGATA | 3594040 |
| 6 | TTATTACCCGTTTTACAGCGTTATG | 3600483 |
| 7 | CTTGTCTTTGCTAAGGTTTTCAATC | 293116 |
| 8* | AAATGTCGGGACAAAAGTGC | 286982 |
| 9 | GCTTGACAGAGTGTAAAACAAAACAT | 3551517 |
| 10* | CTGGGCGAATTCGATGATAC | 3558554 |
| 11 | TATAAGAAAATGGGATTCAAGGTGA | 4263010 |
| 12 | GATGAAAAATCAACAAACAGAAAAGA | 4257133 |
| 13 | GTTAGAGAAAGCACGTTCCTTGTAG | 3743630 |
| 14 | AATGTTCTTCCTTTCTCTTTCGTTT | 3749552 |
| 15 | CCTACTCTCTGTCGAGTAGTGAACTG | 3418263 |
| 16 | TGAATAATGGAGTAACACGTGAAAA | 3424386 |

**Supplemental Table S1.** Sequences of long-range PCR primers used to identify GS. *Adapted from Kothapalli et al. 2005*.*

| Isolate | # Reads spanning 3-5 bridge | # Reads spanning 5’-3 bridge | Bridging read ratio* | Interruption |
| --- | --- | --- | --- | --- |
| T | 208 | 111 | ~2:1 | Mixed GS population |
| EZ T | 87 | 100 | ~1:1 | Single GS population, 1’6’5’35642’7 |
| ISO T | 305 | 165 | ~2:1 | Mixed GS population |
| LAT2 | 189 | 0 | 1:0 | Single GS population, GS21.3 1’35642’7 |

**Supplemental Table 2.** Summary of bridging reads. Number of reads which bridged fragments 3 and 5 (3-5) and fragments 5’ and 3 (5’-3) in the T variants. LAT2 was used as a method control. *Bridging read ratio is given as 3-5:5’-3. A ratio of 2:1 indicates the equal presence of both GS21.3 (1’35642’7) and 1’6’5’35642’7. A ratio of 1:1 indicates the presence of 1’6’5’35642’7 only and a ratio of 1:0 indicates GS21.3 (1’35642’7) only.

**Supplemental Methods**

**PMAxx** **Real-Time PCR Bacterial Viability Test**

A culture of WT was grown overnight at 37 °C, 180 rpm before being OD adjusted to match that of the glycerol stock of T, thus allowing WT to have roughly the same number of cells as that seen in the glycerol stock. To provide controls, this was split into two 1 mL aliquots (~4x10^7^ cells/mL), of which one aliquot was labelled live and the other was subjected to heat shock (95 ◦C for 5 min) and labelled dead. The corresponding live and dead aliquots, prepared in triplicate, were used to make 400 µL samples which contained 0, 25, 50, 75 and 100 % live cells. A sample of glycerol T was also prepared by harvesting the cells within 400 µL and resuspending them in LB supplemented with aro mix. All samples were then stained with PMAxx according to the manufacturer’s protocol. Briefly, 100 µL of 5X PMA Enhancer and the membrane impermeable PMAxx dye (25 µM) was added, before being incubated for 10 min in the dark on a platform rocker. Then, the sample was exposed to light for 15 min to cross-link PMAxx to DNA in non-viable cells with compromised cell membranes. Cells were then harvested by centrifugation at 5,000 g for 10 min and DNA was extracted using GeneJET protocol as per manufacturer’s instructions. 2 µL of extracted DNA was used in qPCRs with 10 µL Forget-Me-Not Master Mix, 0.5 µM *invA* primer mix in a total volume of 22 µL. The qPCR conditions were: pre-incubation at 95 °C for 5 min, amplification for 40 cycles at 95 °C for 5 sec and 60 °C for 30 s, and melt curve performed between 57 and 99 °C for 1 min, with a final hold at 37 °C. The Ct values obtained for live/dead control samples were plotted against live cell numbers, obtained by OD growth curves, and used to determine the number of live cells within the T glycerol stock.

**PCR confirmation of *ΔaroC* in WT derivatives**

To confirm that the picked colonies were derivatives of WT and not contamination, a PCR was performed using primers amplifying *aroC*. WT contains a deletion in aroC and so can be clearly distinguished among other *Salmonella*. The aroC PCR amplified a 1010 bp fragment for the wild type *aroC* and a 360 bp fragment in *ΔaroC* WT mutants. DNA for *aroC* PCR was obtained from single colonies that were lysed in 50 µL nuclease-free water via incubation at 99 °C for 10 min. PCRs were performed on 1 µL of DNA with 1.1X PCR SuperMix (Invitrogen), 0.5 µM forward primer (GACAACTCTTTCGCGTAACC) and 0.5 µM reverse primer (GTGATCCATCAGTACGATCG) in a total volume of 26.25 µL. The PCR conditions were: pre-incubation at 95 °C for 50 sec, amplification for 25-35 cycles at 95 °C for 10 sec, 55 °C for 1 min and 72 °C for 1 min, with a final extension at 72 °C for 1 min.

**Confirmation of GS structure/s in T variants**

As assemblies of long-read sequences of T variants were unable to be resolved due to the potential presence of mixed GS populations of 1’6’5’35642’7 and 1’35642’7, we searched the filtered reads for those which spanned fragments 3 and 5 and fragments 5’ and 3. Reads were searched for three different 25 base pairs using seqkit fish command which is specifically designed to look for short sequences in long read sequences, whilst accounting for potential errors and the reverse complement. These sections were unique, as only located once in the parent WT hybrid assembly. These three sections, which are described here in relation to their positions in the parental genome structure GS2.66 (17’35642’), are as follows: 1) 25 bp section in fragment 5, 100 bp away from fragment 3 (AATGATGTATCGCAGATTTCTGCCT); 2) 25 bp section in fragment 3, 400 bp away from fragment 5 (AAATAAGCAAATTGCCGTTATTGCA) and 3) 25 bp section in fragment 3, 300 bp from fragment 7 (AAAAATCGCCACTTTGCGCAGGAAT). Results of these separate searches were then compared and used to find reads which bridged fragments 5 and 3 (searches 1 and 2, respectively) and fragments 5’ and 3 (searches 1 and 3, respectively).
